# Build-up of serial dependence in color working memory

**DOI:** 10.1101/503185

**Authors:** João Barbosa, Albert Compte

**Affiliations:** Institut d’Investigacions Biomèdiques August Pi i Sunyer (IDIBAPS), Barcelona, Spain

## Abstract

Serial dependence, how recent experiences bias our current estimations, has been described experimentally during delayed-estimation of many different visual features, with subjects tending to make estimates biased towards previous ones. It has been proposed that these attractive biases help perception stabilization in the face of correlated natural scene statistics as an adaptive mechanism, although this remains mostly theoretical. Color, which is strongly correlated in natural scenes, has never been studied with regard to its serial dependencies. Here, we found significant serial dependence in 7 out of 8 datasets with behavioral data of humans (total n=760) performing delayed-estimation of color with uncorrelated sequential stimuli. Moreover, serial dependence strength built up through the experimental session, suggesting metaplastic mechanisms operating at a slower time scale than previously proposed (e.g. short-term synaptic facilitation). Because, in contrast with natural scenes, stimuli were temporally uncorrelated, this build-up casts doubt on serial dependencies being an ongoing adaptation to the stable statistics of the environment.

## Introduction

Our perception depends on past experiences [1]. Serial dependence - how our current estimates are biased towards previous ones - has been described experimentally using many different paradigms [2–18]. In particular, paradigms including delayed-estimations of different visual features [2], such as orientation [9,11,15], numerosity [19], location [3,20,21], facial identity [22] or body size [13]. It has been speculated that these ubiquitous attractive biases are a consequence of the world’s tendency to be stable, and have the functional role of averaging internal noise [2,11,14,23]. Some have further argued that serial dependence is of adaptive nature, changing its strength depending on the stimuli statistics [2,11,14,16,23]. Color, which is strongly correlated in natural scenes [24], has never been studied with regard to its serial dependencies, possibly due to its strong systematic biases [25–27]. Similar to other perceptual biases for other visual features [28,29], these systematic color biases adapt to stimulus statistics in the course of one experiment [27]. This suggests that typical perceptual bias adaptations occur in time scales of minutes to hours. Slow adaptation of serial dependence, however, has never been characterized. If serial biases are also subject to adaptation with a similar time scale, when exposed to long sessions with uncorrelated stimulus statistics they should decrease or, in case of not being adaptive, they should remain stable. In fact, a recent study supports the latter hypothesis: in an auditory working memory task, with sound frequencies sampled from uniform, gaussian or bimodal distributions, serial biases were not affected by the stimulus distribution [18]. Here, we address serial dependence in delayed-estimation color tasks, controlling for systematic biases and - contrary to our hypothesis - we characterize for the first time a slower dynamics of increasing serial dependence through the experimental session, despite uncorrelated stimulus statistics.

## Results

### Serial dependence in color working memory

We analysed 8 datasets that are freely available online (see Methods), with a total of n=760 subjects performing variations of the same, delayed estimation of color task (Figure 1a, Table S1). We found that, across experiments, the subjects’ reports were attracted to the previous target color for relative distances between previous and current trial target color of up to 90° in all experiments. Significant serial dependence occurred in all individual datasets for relative distances of up to 90° (Cam-Can, p=1.07e-09; Van der Berg I, p=0.006; Van der Berg II, p=9.66e-06; Oberauer & Li, p=0.007; Foster et al I, p=0.002, Foster et al II, p=4.35e-08; Bays et al, p=0.003), except for the dataset collected by Souza et al [30] (p=0.14).

**Figure 1.**
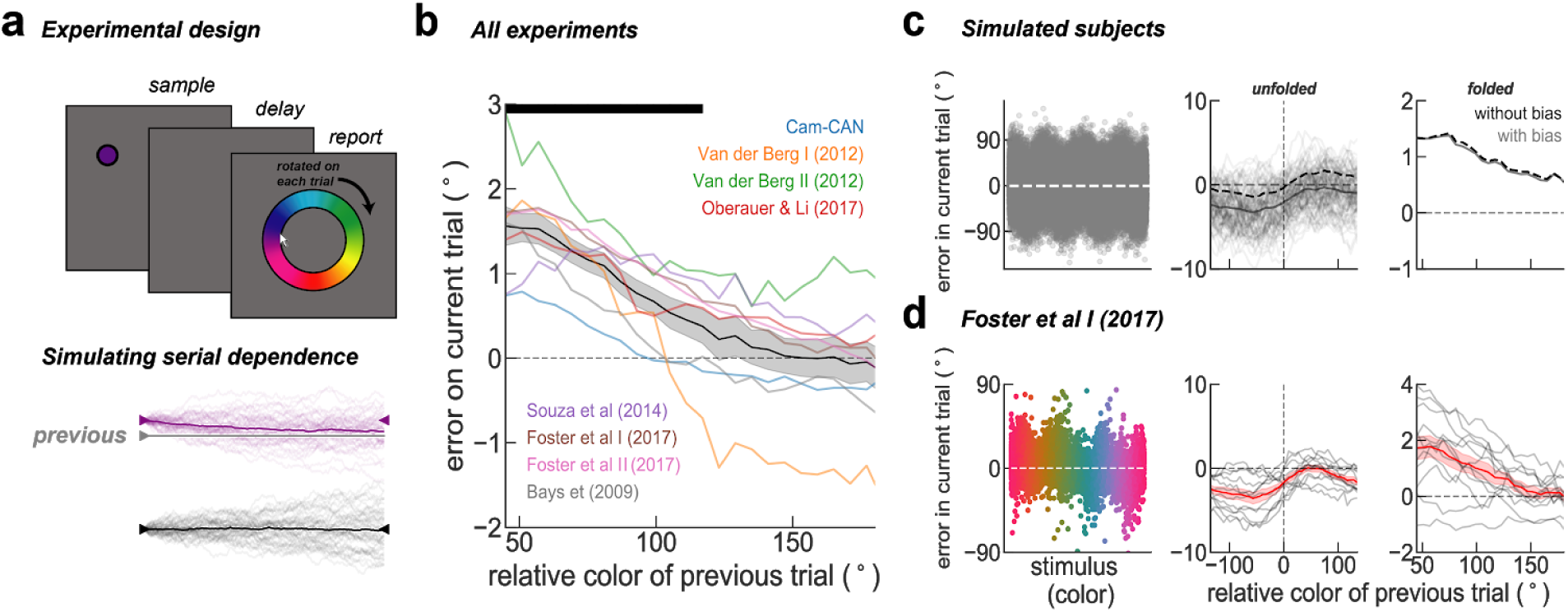
serial dependence in color. a) Top, experimental design. All experiments were variations of a delayed-estimation of color task as in ref. [31], differing mostly on set size and number of trials (Table S1). Subjects reported on a color wheel rotated by a random angle in each trial. Bottom, serial dependence was simulated as a drift towards the previous trial trace in a diffusion process. In purple, 50 trials with a stimulus feature (purple triangle) close to previous trial trace (gray) and in black, 50 far trials. Thick lines represent the averages of each condition, which are attracted to previous trial stimulus for trials that are close by. b) Serial bias in the delayed-estimation of color task for all datasets. We found significant serial dependence relative to previous report in all datasets (p=0.0003, t-test), except for the dataset collected by Souza et al [30] (p=0.14, t-test). c) Left, error to target stimulus reveals systematic biases on simulated trials. Middle, serial dependence calculated separately for trials simulated with and without systematic bias. Right, folded version of serial dependence removes all systematic biases without any additional preprocessing. d), same as c) for trials of Foster et al I [32].

### Folding serial bias curve removes systematic biases

Delayed-response reports are subject to systematic biases, which are particularly strong in the case of color [26]. It has been argued that it is necessary to model and remove the systematic bias prior to estimating serial dependence [3,9,21]. Here, we applied a model-free strategy that corrects serial dependence by “folding” the serial bias plot (Figure 1b). We tested this method in surrogate data obtained using a computational modeling approach. We simulated each delay of two consecutive trials as a diffusing memory trace [33] using a simple random walk simulation (see Methods for details). On top of independent Gaussian errors responsible for diffusion, we added serial dependence as another source of error that accumulated incrementally at each time step, and two other sources of distortion (see Methods for details): 1) systematic biases derived from inhomogeneities of the task space, and 2) systematic rotational biases (e.g. a constant clockwise error [3,9,15]). Figure 1c shows the effect of these systematic biases on serial bias estimation. We simulated n=1000 trials and n=100 subjects with (gray) and without (dashed black) systematic biases. As previously reported [9], we found that systematic biases shift the serial bias function to negative values. This shift precludes the correct identification of attractive and repulsive serial bias regimes, and complicates comparison across subjects. We found that a simple processing of the data allowed for a model-free correction of systematic-bias-induced shifts: we “folded” the serial bias curve by collapsing all negative distances between consecutive targets on positive values, while also inverting for these trials the sign of the behavioral error (*folded error,* see Methods). This method effectively removes all systematic biases introduced in simulated trials (Figure 1c, right). For illustration purposes, we show the application of this method in one dataset (Foster et al I [32]) with similar systematic biases (Figure 1d, left), that led to a shifted serial bias function (Figure 1d, middle) and finally a *folded* version, without systematic biases (Figure 1e, right). See Figure S1 for same analyses on each experimental dataset. Thus, our simulation approach validated the folding approach to correct for systematic biases in serial bias estimations, and allow for across-subject comparisons (Figure 1b).

### Color serial dependence builds up in the course of an experimental session

Serial dependence, some argue, reflects the world’s tendency to be stable [2,14,23]. The reasoning is that because similar stimuli usually elicit similar behavior, the brain would incorporate mechanisms to exploit these patterns [23]. Along these lines, a recent study has shown that systematic biases in color working memory change in the course of an experimental session to adapt to stimulus statistics [27], arguing that systematic biases seen in delayed-estimation of color reflect real-world statistics. If similar adaptive plasticity operated for serial dependence in the time scale of the experimental session, we would expect to see a reduction of serial biases as one is exposed to a sequence of uncorrelated stimuli. To our knowledge, the stability of serial dependence within an experimental session is yet to be characterized. To address this question, we divided each session in two halves and computed serial dependence relative to the previous trial report for each subject and experiment in each of these two halves (Figure S2). When averaging all experiments together, we found that, contrary to our hypothesis, there was stronger serial dependence in the second half than in the first half of the experimental session (Figure 2a). To further characterize this serial dependence build-up, we used the folded error on each trial as a scalar measuring the evolution of serial dependence in the course of the session (see Methods). Figure 2b illustrates this analysis using a sliding window of 75 trials for Cam-Can [34,35] dataset and of 200 trials for the Foster et al I dataset [32], both showing a clear increase of serial dependence as the session progressed. To test this effect across subjects and experiments, we obtained the regression slope of the folded error as a function of trial number. We computed this slope for each subject and for all experiments. The data shows that serial dependence build-up was positive for all but 1 dataset (Bays et al [36]), significant for 2 datasets individually (Cam-Can [34,35], p=0.008 and Foster et al I [32], p=0.0002, t-test) and for all combined (p=0.043, t-test on the 8 averages across experiments or combining subjects from all experiments, p=0.006).

**Figure 2.**
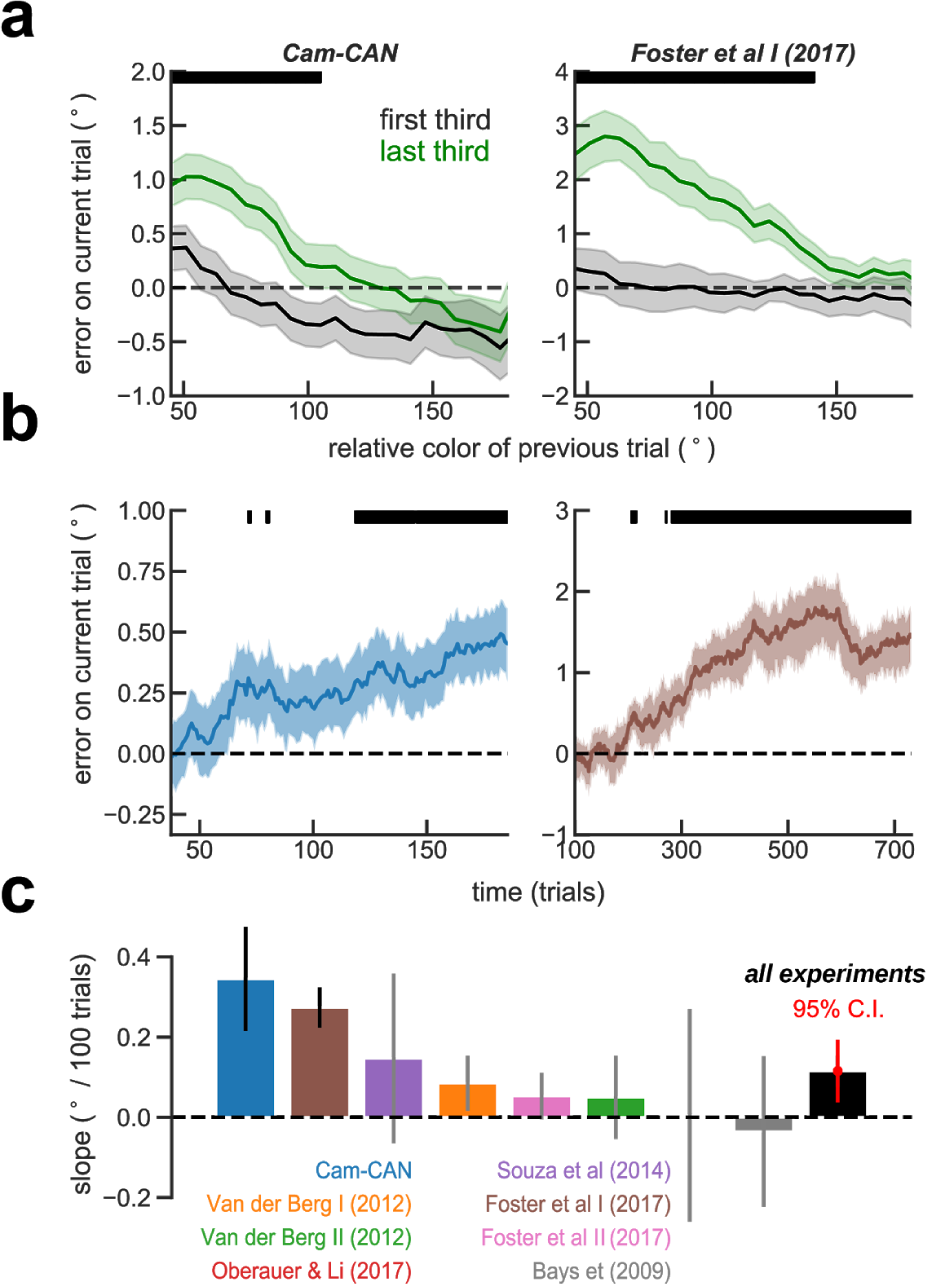
Serial bias builds up during a session. a) Serial biases computed using first third (black) and second third (green) of the trials for two example experiments: Cam-Can and Foster et al I [32]. Black bars on the top mark where curves are significantly different, p<0.05, permutation test. b) Both experiments show a significant increase in serial dependence through the session computed with a sliding window of 75 trials for Cam-Can [34,35] and 200 for Foster et al I [32]. c) For each subject, we computed the slope of serial dependence over the course of the session (without averaging). We found that serial-bias build-up was significant in two experiments (marked with black error-bars: Cam-Can, p=0.008; Foster et al I, p=0.0002). Error-bars were calculated from bootstrap distributions and unless stated otherwise, are standard errors of the mean.

We then tested if this serial bias build-up was related with subjects getting familiar with the task, in which case one would expect to see an improvement in performance through the session, or related to subjects feeling tired, which should be reflected in worsening of performance. To this end, we calculated the fraction of guesses as a proxy of tiredness or engagement. We classified as guesses those trials with error > 90° in independent windows of 20 trials. Importantly, these trials were excluded from all the other analyses (above). We then computed the slope of change of the fraction of guesses in the course of the sessions for each participant. We found a decrease in guess rate through the session in 2 datasets (CamCan, p=7.1e-20 and Van der Berg I, p=0.008) and an increase in another 2 datasets (Souza et al, p=0.02 and Bays et al, p=1.1e-06). However, these trends did not correlate with serial bias build-up for any dataset independently (p>0.35, linear regression, Figure S3a), combining subjects from all datasets (p>0.35, linear regression) or averaging across experiments (p>0.2, linear regression, Figure S3c). Also, we checked the evolution of performance by measuring the mean squared error through the session. Serial bias build-up was also not correlated with the subjects’ squared error (excluding guess trials) trend during the session (Figure S3). These control analyses show that serial bias build-up during experimental sessions was not associated with trends in performance dynamics as a result of subjects getting familiar with the task or tiredness. Together, these results show that serial dependence is not stable on the time scale of one experimental session, as previously assumed, and it also discards a mechanism that adapts to stimulus statistics. Instead, our result suggests the involvement of slowly accumulating plastic mechanisms in serial dependence of color delayed-estimations.

## Discussion

We provide the first evidence of serial dependence in color working memory. Serial dependence had been characterized with great detail for other visual features [2], and in particular in spatial working memory [3,20,21]. Several common features of color and spatial working memory suggest that serial dependence could also be similar in color: simultaneously memorized stimuli interfere attractively when presented at close distances [37,38], and memory precision decreases with memory period duration [3,39,40]. These commonalities are in contrast with the differences of neural representations. While spatial representations consolidate early in the visual pathway [41], complex transformations in color representations occur as color information travels from the photoreceptors in the retina, to visual cortex, and into association cortex [42]. The fact that serial dependence is similar for color and spatial working memory thus suggests that it depends on inter-trial interferences that occur at processing stages with representational maps equally distant from the corresponding perceptual map, and this points at higher color processing stages. A candidate region for this is the inferotemporal (IT) cortex, where continuous neuronal representations of color of circular shape on the two perceptual cardinal axes (yellowish-bluish and greenish-reddish axis) have been found [43,44].

The analogy of color and angular location neural representations motivated us to simulate color working memory similarly to spatial working memory of angular locations [45]. We simulated the angular memory trace in the memory period as a diffusion process [33] with a drift toward the previous trial memory trace that introduces serial dependence [46]. We used this model to test the concerns about the impact of systematic biases in the estimation of serial dependence. This is a general concern that has been raised for other visual features [3,9,20,21], but in the case of color it may be particularly important for the marked perceptual systematic biases that have been reported [26]. We therefore incorporated strong systematic biases in the reports of our model simulations, we tested the impact on the estimation of serial dependence and we developed new analysis strategies to address this. One typical strategy for systematic bias removal is to low-pass filter the responses as a function of stimulus feature [3,9,20,21]. This approach depends on parameters that are often subjectively decided (e.g. size of sliding window). In addition, removing systematic biases incorrectly, for example when subjects do not have systematic biases, can introduce extra biases in otherwise clean data (Figure S4). We showed that by folding the serial dependence function, one can reduce the impact of systematic biases on serial dependence without adding biases in unbiased data and without specifying arbitrary parameter values. We therefore conclude that this analysis allows a more robust estimation of serial dependence in behavioral studies.

Theoretical models have proposed that short-term subthreshold mechanisms in inter-trial intervals underlie serial dependence in delayed-estimation of location [46–48]. In this class of models, neural activity in previous trial’s mnemonic representations engage plasticity mechanisms that leave a selective trace in the network’s synapses. This trace interferes with neural activations in the next trial by biasing the neural representation of the new stimulus towards the previous memorized location. These dynamics explain most experimental findings of serial dependence in spatial working memory [46], and our simplified modeling approach is consistent with this mechanistic substrate [46]. However, our finding that attractive serial biases build up in the course of an experimental session is not explained by these models. Indeed, the short-term synaptic plasticity mechanisms invoked so far operate in time scales of a few seconds, much shorter than the time scale of the experimental session. Our results reveal that additional mechanisms, accumulating in a time scale of 10’s of minutes or hours, are also responsible for the instantiation of serial dependencies in delayed-estimation of color tasks. Possible mechanisms are changes in plasticity efficacy itself (i.e. “metaplasticity”), modulating synaptic release probability over the experimental session. We tentatively speculate that habit-related endocannabinoid modulation of synaptic release [47,49] could mediate serial dependence build-up as a non-adaptive result of task habituation.

The build-up of serial dependencies during an experimental session has further implications for how we interpret their functional role. If serial dependence was an adaptation to exploit the world’s tendency to remain stable [23], and this adaptation could occur in the time scale of hour fractions (as recently shown for systematic biases in delayed-estimation of color tasks [27]), memorizing a sequence of uncorrelated stimuli should decrease serial dependence in the course of an experimental session. Alternatively, if hard-wired mechanisms underlie serial dependence, we wouldn’t expect any change. Instead, our results show that serial dependence builds up, suggesting that it does not respond to an active adaptation to the statistics of visual stimuli in the environment (at least in the time scale of hours) but instead may reflect a plasticity mechanism driven by repeated selective neuronal activations in the circuit.

## Methods

### Participants and design

We analysed 8 datasets that are freely available online (Table S1), with a total of n=760 subjects performing variations of the same, delayed-estimation color task (Figure 1a). We will briefly describe the general experiment and Table S1 summarizes the specifics of each task, for detailed descriptions please refer to the original studies [30,32,36,50,51]. On each trial, a set of colored stimuli (varying from 1 to 8 stimuli) were briefly shown. After a delay period of roughly 1 second (see Table S1 for details), during which stimuli were no longer visible, subjects had to report the target color of a cued location. These color reports correspond to angles (i.e. degrees) on a color wheel rotated by a random amount on every trial, to avoid a spatial memory strategy.

### Serial dependence analysis

As in previous studies (e.g. ref. [11]), serial dependence was measured by calculating the mean error as a function of distance between current and previous target. Positive (negative) errors for positive (negative) distances are interpreted as attraction towards the previous target while negative (positive) errors for positive (negative) distances as repulsion. Most of the studies, except for Foster et al [32], were multi-item working memory. On trials with more than one stimulus, we use the target stimulus as reference.

#### Folded errors

We corrected for the effect of systematic biases in color estimation [26] by computing a folded version of typical plot (e.g. ref. [11], see Discussion) as follows. We computed “folded” error (multiplied by the sign of the target-to-target distance) as a function of absolute distance between targets.

#### Serial dependence build-up

Serial dependence build-up was measured by splitting each session’s trials in three equal consecutive thirds and computing serial dependence separately. Serial dependence build-up was also measured by using the whole session data. For the whole session data analyses, we computed the folded error on each trial (by multiplying each error with the sign of the target-to-target distance, see above). For each session, we fitted a regression slope of folded error against trial number and averaged across sessions, keeping one slope per subject and experiment.

### Simulating consecutive trials

We simulated the memory trace in trial *k* as a random walk, 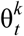(t=0..T, with T the duration of the delay period). Each random walk (Eq. 1) started at its corresponding stimulus location, 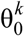. On top of independent Gaussian errors ξ_*t*_ responsible for diffusion (controlled by the parameter *σ=0.006* deg), we added serial dependence as one more source of error that accumulated incrementally at each time step. This increment was obtained from a derivative-of-Gaussian (DoG) function of the distance between the instantaneous memory trace and the previous trial’s stimulus (i.e. starting or ending location of the previous trial simulated trace 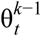, with t=0 or t=T, respectively) (Eq. 2, [9,11,15,22]) The DoG curve was defined by parameters *a = 0.09* deg, 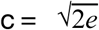, and *w=0.007* deg^−1^ (Eq. 2), similar to [9].

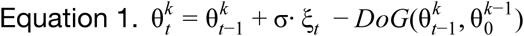

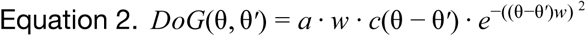

The *response* in each simulated trial was then derived from the last point of the memory trace 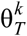, by applying two sources of distortion: 1) systematic biases derived from inhomogeneities of the perceptual space were simulated as 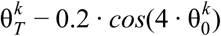, with constants chosen to fit the data by visual inspection (Fig. 2c,d, left); 2) systematic rotational biases (e.g. a constant clockwise error [3,9,15]) were simulated by adding a constant arbitrary angle 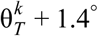

### Statistical tests

All error-bars and confidence intervals were computed from bootstrapped distributions. Significance was assessed with 1-sample *t*-tests or permutation tests at P=0.05.

## Acknowledgments

This work was funded by the Ministry of Economy and Competitiveness of Spain, and the European Regional Development Fund (Refs: BFU2015-65318-R, RTI2018-094190-B-I00), by the Secretaria d’Universitats i Recerca del Departament d’Economia i Coneixement de la Generalitat de Catalunya (Ref: SGR17-1565), and by the CERCA Programme/Generalitat de Catalunya. JB was supported by the Spanish Ministry of Economy and Competitiveness (FPI program). This work was developed at the building Centro Esther Koplowitz, Barcelona. Data collection and sharing for this project was provided by the Cambridge Centre for Ageing and Neuroscience (CamCAN). CamCAN funding was provided by the UK Biotechnology and Biological Sciences Research Council (grant number BB/H008217/1), together with support from the UK Medical Research Council and University of Cambridge, UK. We thank the generous sharing of behavioral data that facilitated this work, especially to Paul Bays, Joshua Foster, Klaus Oberauer, Alessandra Souza, Ronald van den Berg and Daniel Mitchell for their specific supportive interactions. We also thank Daniel Linares, Heike Stein and Matthias Fritsche for critically reading early versions of the manuscript.

## Author contributions

JB and AC designed the study, JB analysed the data and JB and AC wrote the manuscript.

## Data availability

Each dataset is available from the original publications [30,32,36,50,51] and all code used for the analyses will be available at https://github.com/patra/serial_color after publication.

