## Supplementary Material for "Build-up of serial dependence in color working memory"

### Supplementary Figures

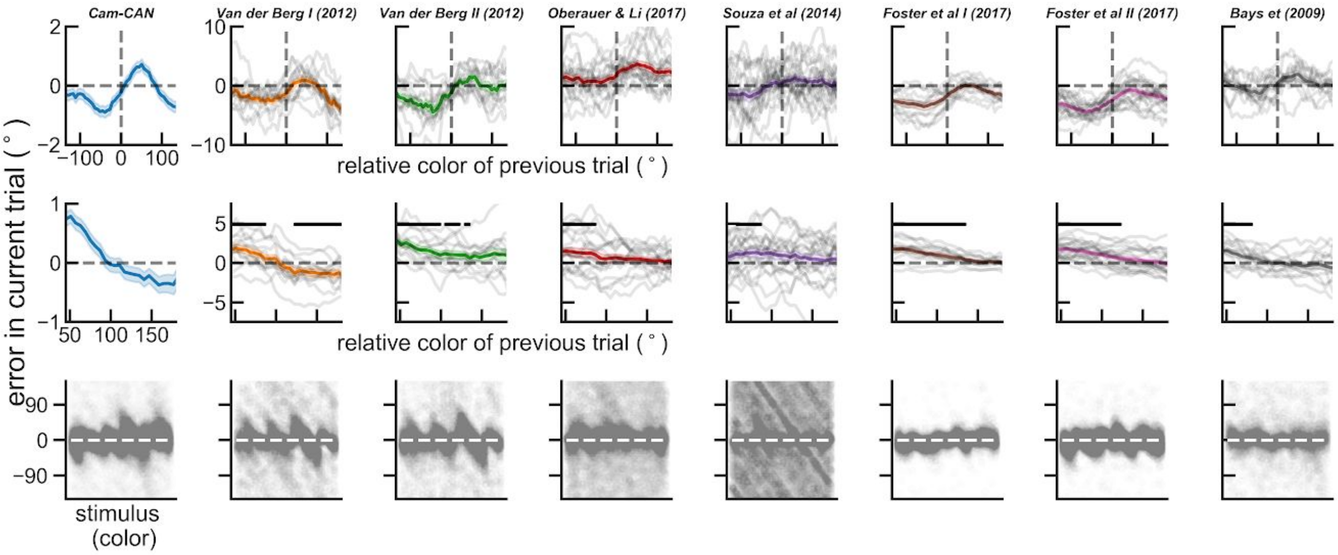


**Figure S1. Serial dependence in color (folded and unfolded) for each experiment.** Serial dependence in color for each experiment, unfolded at the top and folded in the middle. At the bottom, systematic error for each experiment. For the Cam-Can dataset, we only plotted the systematic error from 100 subjects for the sake of clarity.

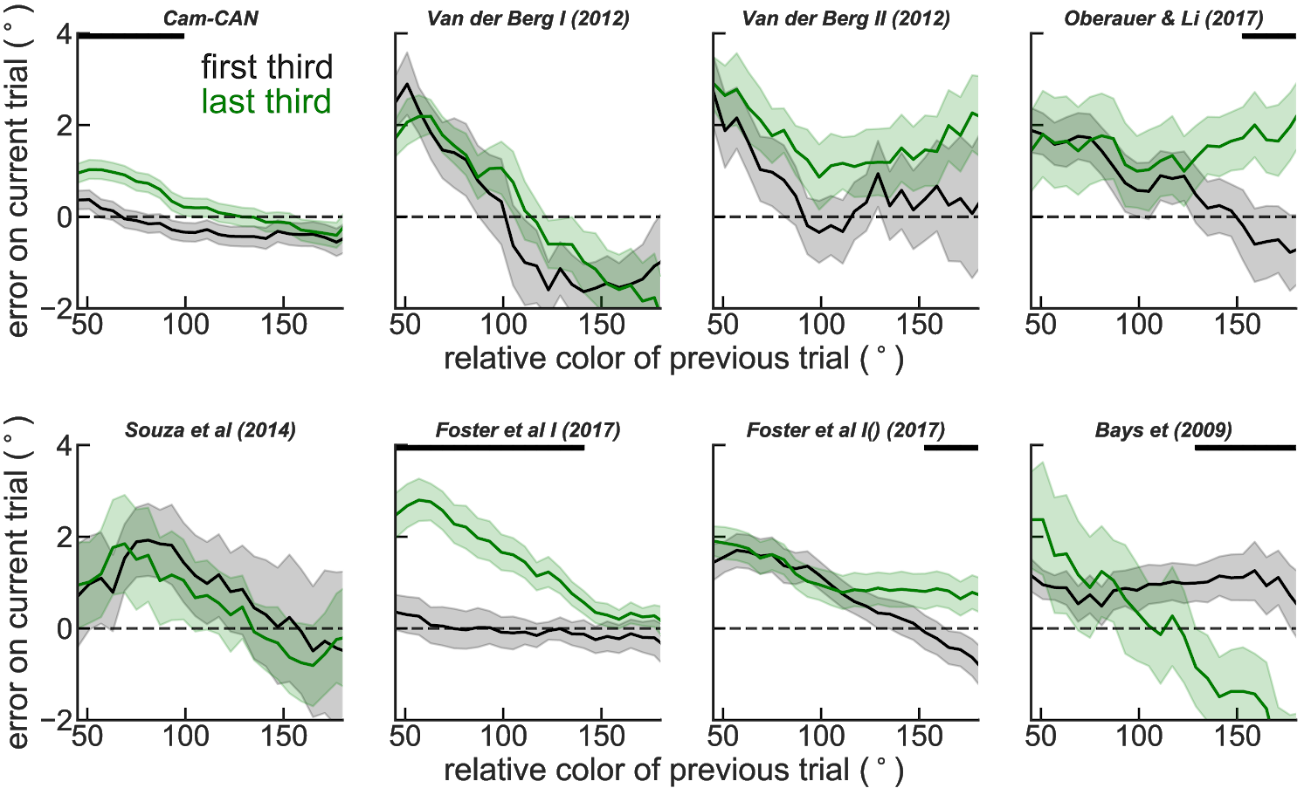


**Figure S2. Serial dependence calculated using first and second half of each session.** All error bars are bootstrapped SEM and black bars mark where the two conditions are significantly different ($p<0.05$, permutation test).

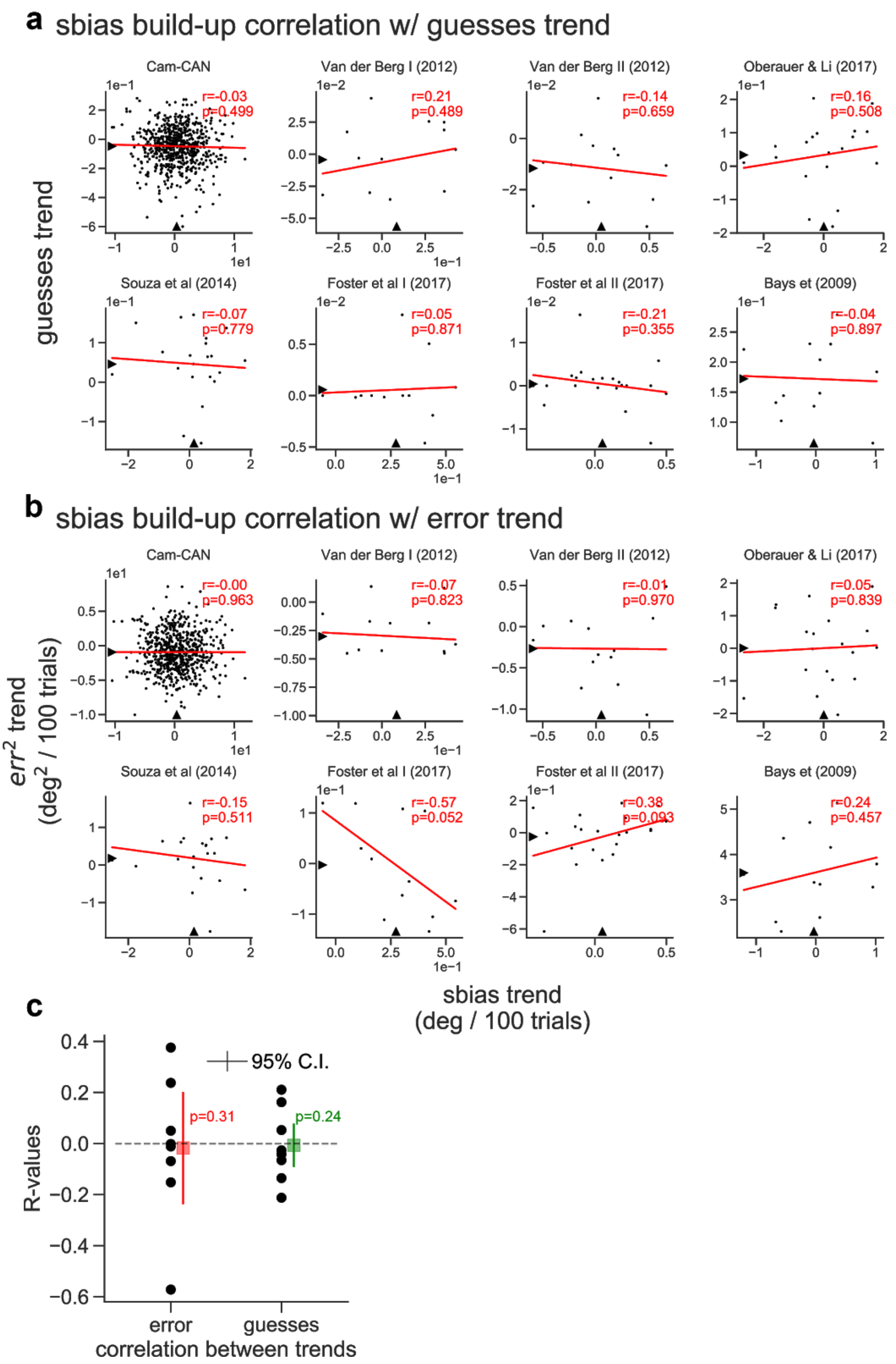


**Figure S3.** Serial bias build-up is independent of tiredness or task familiarity. Serial biases build-up is not correlated with trends in performance dynamics as measured by a) guesses or b) mean squared error for any dataset independently (a, b) or averaging across experiments (c).

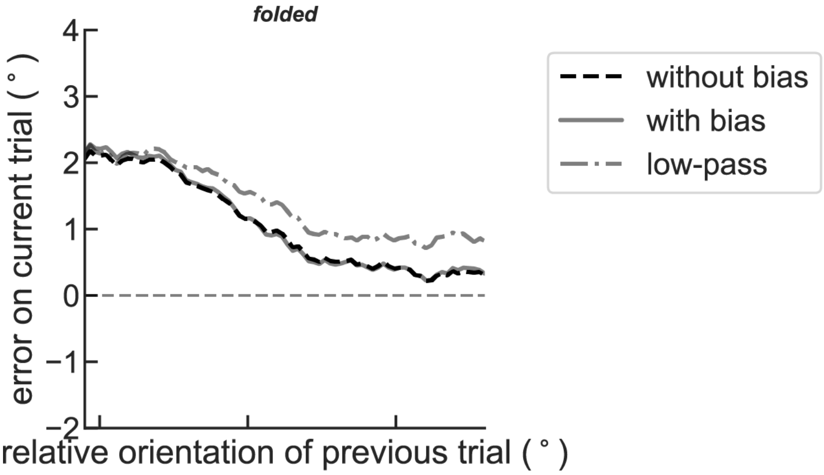


**Figure S4.** Replication of Figure 1c. A folded version of serial dependence removes all systematic biases without any additional preprocessing (compare curve “with bias” with curve “without bias”). In addition, here we also plot serial dependence from the data “with bias” when removing systematic biases with a low-pass filter similar to refs. [1–3] prior to folding. Notice that the “low-pass” curve introduces biases in the estimation of serial dependence (compare to “without bias” curve).

| *Dataset* | *Set size* | *Subjects* | *Trials* | *Delay* | *Observations* |
|---|---|---|---|---|---|
| CamCan data set<br>Taylor et al (2017) [4]<br>Shafto et al (2014) [5] | 1-4 | 649 | 224 | 0.9 s | Stimuli: circle of diameter 1.77 (dva), positions selected at random from 8 equally spaced points at an eccentricity of 4.5. 360 colors. CIE L,a,b radius of 53 and center (64,10,10). Half of the trials had non-targets probing feature revealed. These data was obtained from the CamCAN repository, available at: www.mrc-cbu.cam.ac.uk/datasets/camcan/ |
| Experiment 1 of<br>Oberauer & Li (2017) [6] | 1-8 | 19 | 400 x 2 | 1 s | Stimuli: colored squares of 1.25º at viewing distance of 50cm. 360 different colors on a color wheel: CIE L,a,b = (70,20,30). |
| Experiment 1 (I) of<br>Foster et al (2017) [7] | 1 | 12 | ~960 | 1.2 s | Stimuli: circle, 1.6º diameter, centered at 3.8º at viewing distance of 100cm. 360 colors. Color wheel in Figure 1 |
| Experiment 2a (II) of<br>Foster et al (2017) [7] | 1 | 21 | ~960 | 1.15 s | Stimuli as I, plus: During presentation, a distractor with different shape and color was present at another location. |
| Experiment 1 (I) of<br>Van den Berg et al (2012) [8] | 1-8 | 13 | 288 x 3 | 1 s | Stimuli: circle, 2º diameter, centered at 4.5º at viewing distance of 60cm. 180 different colors on a color wheel: CIE 1979, L,a,b = (70,10,10). |
| Experiment 3 (II) of<br>Van den Berg et al (2012) [8] | 1-8 | 13 | 288 x 3 | 1 s | Same as I, but report done by scrolling through all possible colors (drawn uniformly and independently from the wheel). |
| Experiment 2 of<br>Souza et al (2014) [9] | 1-8 | 21 | 496 x 2 | 1 s | Stimuli: circles of 1.1 cm diameter at 5.5cm from fixation. 360 different colors samples from hue dimension of HSL (saturation=1, lightness=.5). Condition 1: color wheel present during delay Condition 2: Last 1 sec of - sec delay was location cued. |
| Bays et al (2009) [10] | 1-8<br>(in blocks) | 12 | 600 | 0.9 s | Stimuli: 2x2º patches at viewing distance of 57cm. 180 different colors samples from CIE L,a,b=(70,20,38). Presentation duration varied: 0.1, 0.5, or 2 s in a block design. This could potentially confound build-up analysis. |

**Table S1. Experimental details of each dataset.** With the exception of Foster et al I & II [7], all datasets have varying set size.
